# A gene-augmentation framework for selected early-stage autosomal recessive retinitis pigmentosa genotypes

**DOI:** 10.64898/2026.07.25.740701

**Authors:** Peimin Ma, Xiaomei Sun, Shuya Xu, Mengxian Yang, Chen Gao, Xin Chen, Li Gong, Wen Zeng, John J. Renger, Yunlu Xue

## Abstract

Many recessive inherited retinal dystrophies are, in principle, amenable to gene augmentation, yet only one such therapy has received regulatory approval. We sought to identify potentially tractable autosomal recessive retinitis pigmentosa (RP) genotypes that might be addressed using publicly described adeno-associated virus (AAV) components. More than 100 RP-associated genes were prioritized according to cellular expression, coding-sequence size, and the availability of cell type-directed promoters, yielding 19 candidate genes expressed predominantly in rods and/or retinal pigment epithelium (RPE). In rhesus monkey eyes, the human RHO and BEST1 promoters supported rod- and RPE-enriched expression, respectively, whereas the tested GFAP and RLBP1 constructs were limited by undetectable Müller glial expression or RPE abnormalities under the tested exposure conditions. We selected PDE6B as a proof-of-concept gene and evaluated AAV8-RHO-PDE6B after neonatal subretinal delivery. The vector preserved outer nuclear layer structure, electroretinographic responses, and visually guided behavior in rd1 and rd10 mice, with effects observed through 6 months. Higher tested doses were associated with retinal abnormalities, whereas fewer detectable abnormalities were observed at the lower doses through day 56 in rhesus monkey eyes. Together, these findings define a preclinical framework for developing gene-augmentation vectors for selected early-stage autosomal recessive RP genotypes.

## Introduction

Retinitis pigmentosa (RP), one of the most common inherited retinal degenerations (IRD), affects approximately 1 in 3,000–4,000 people worldwide (Daiger et al., 2007; Hartong et al., 2006). The disease typically begins with rod dysfunction and night blindness (an early-stage manifestation), progresses to loss of peripheral cone-mediated vision and can culminate in severe central vision impairment. More than 100 genes have been associated with RP (https://retnet.org/summaries#a-genes), with autosomal recessive, autosomal dominant, and X-linked patterns of inheritance. Many RP genes are expressed directly in rods, whereas some others function in neighboring retinal pigment epithelium (RPE) or Müller glia (Lem et al., 1999; Pittler and Baehr, 1991; Redmond et al., 1998; Saari et al., 2001). Although some RP genes are also expressed in cones, the mutation(s) may not affect cones directly, as cone degeneration non-autonomously follows rod loss in RP (Carter-Dawson et al., 1978; Mohand-Said et al., 1998; Punzo et al., 2009) (**Figure 1A**), providing a rationale for interventions that preserve rods before extensive secondary cone loss has occurred (Chen and Cepko, 2009; Pang et al., 2008; Scalabrino et al., 2023).

**Figure 1.**
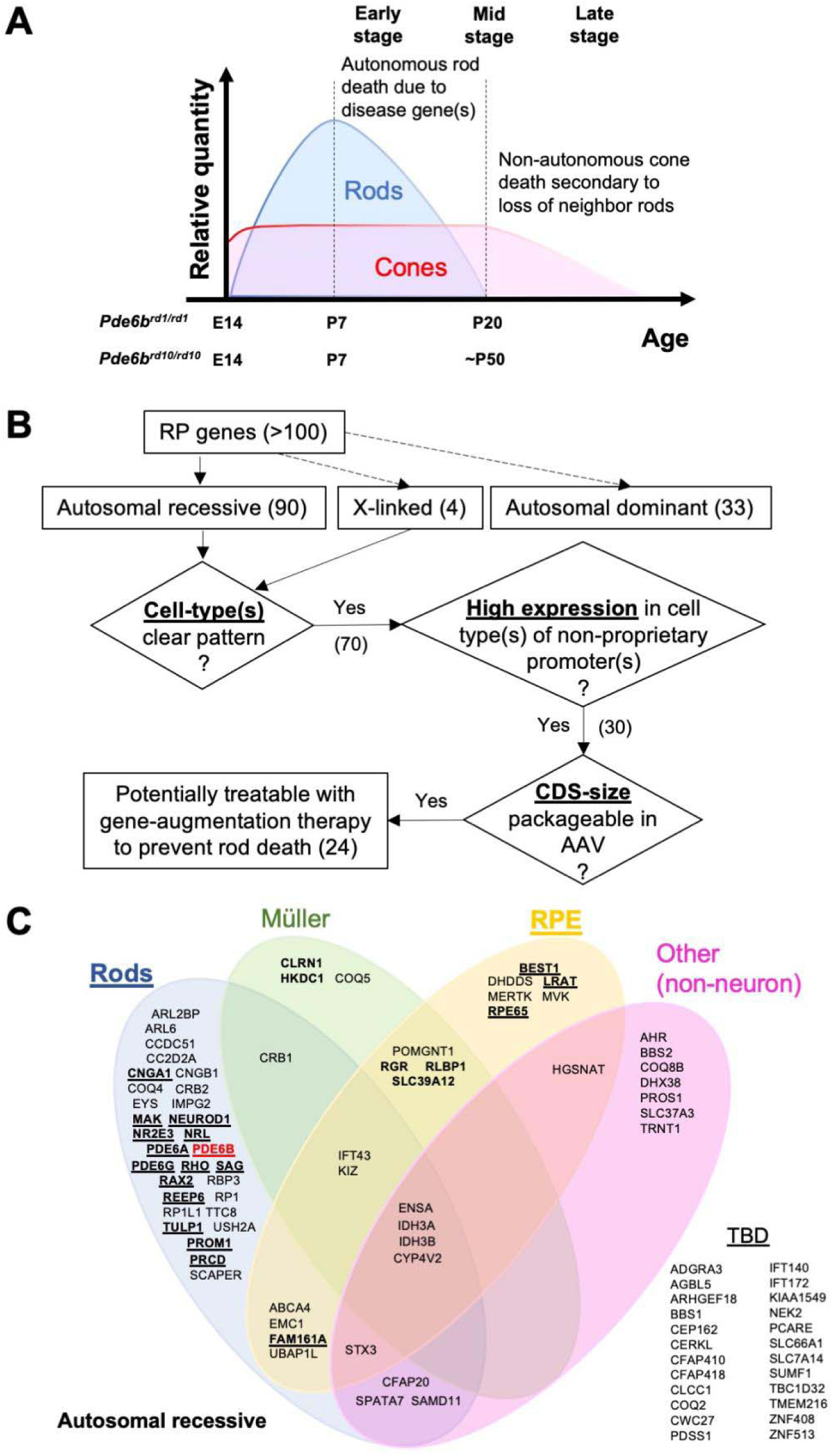
Prioritization of autosomal recessive RP genes for gene-augmentation therapy. **(A)** Schematic of photoreceptor loss in the peripheral retina during RP. Degeneration kinetics are also shown for the rd1 and rd10 mouse strains, both of which carry Pde6b mutations. **(B)** Flowchart of RP gene classification by inheritance. Numbers of genes are shown in parentheses. **(C)** Venn diagram of autosomal recessive RP genes according to cellular expression. Bold, 24 potentially tractable genes compatible with publicly described rod-, RPE-, or Müller-glia-directed promoters reported in mice. Underlined, 19 potentially tractable genes compatible with a non-proprietary AAV8 capsid and rod- or RPE-specific promoters supported by exploratory expression studies in rhesus monkeys (Figures 2 and **S2**). Red, PDE6B, the proof-of-concept gene for the platform.

Gene augmentation is a direct therapeutic strategy for recessively inherited diseases, delivering a functional coding sequence to the cell type in which the affected gene normally acts. Recombinant adeno-associated virus (AAV) is well suited to ocular delivery because it can support sustained transgene expression and has an established clinical record, although its packaging capacity limits the genes that can be accommodated. Voretigene neparvovec, an AAV2-based RPE65 augmentation therapy, demonstrated that subretinal delivery can improve visual function in patients with bi-allelic RPE65-associated retinal dystrophy (Russell et al., 2017). However, it remains the only gene therapy for an IRD. Clinical programs for other genotypes, including RPGR-associated X-linked RP, have encountered safety, efficacy, development, or financing barriers (Lam et al., 2024; Michaelides et al., 2024).

The diversity of RP-associated genes creates two translational challenges. First, an augmentation cassette must fit within the AAV genome, excluding genes that otherwise may be biologically amenable to replacement. Second, a conventional one-product-at-a-time development strategy may be difficult to sustain for the small patient populations associated with individual genotypes. A platform strategy based on shared capsids, regulatory elements, manufacturing processes, and preclinical assays could reduce duplicated development work, but each selected transgene should be individually validated for compatibility by demonstrating efficacy and safety at least in a small animal disease model.

Here, we screened identified RP-associated genes using human ocular cell-type expression patterns, coding-sequence size, and the availability of publicly described cell type-directed promoters. We then tested candidate rod-, RPE-, and Müller glia-directed promoters in rhesus monkey eyes and identified 19 autosomal recessive genes provisionally compatible with the rod-and RPE-directed expression systems supported by these exploratory studies. As a proof of concept, we evaluated an AAV8-RHO-PDE6B-W3SL-TLR9io2 vector in dose-ranging studies in mice and rhesus monkeys and in efficacy studies in the rd1 and rd10 mouse models. This work defines a preclinical experimental approach for a shared gene-augmentation framework for selected early-stage autosomal recessive RP genotypes.

## Results

### A cell-type and coding-sequence screen identifies candidate genes for AAV-mediated augmentation

We first established a set of RP genes that could be addressed using conventional single-AAV gene augmentation. RetNet listed 90 autosomal recessive, 33 autosomal dominant, and 4 X-linked RP-associated genes as of December 2025 (**Figure 1B**). We examined their expression across human retinal and ocular cell types using two published single-cell RNA-sequencing datasets (Voigt et al., 2019a; Voigt et al., 2019b), and cross-referenced the assignments against two additional datasets (Liang et al., 2019; Lukowski et al., 2019). Relative expression was manually adjudicated on a six-level scale ranging from “+++++” to undetected “−” by two scorers, who independently scored and agreed on the final presentation, based on the cell-type percentage of expressing and number of transcripts (**Supplementary Lists 1 and 2**).

Genes were grouped according to predominant expression in rods, Müller glia, RPE, or other non-neuronal ocular cells (**Figures 1C** and **S1**). To reduce the influence of low-level signals and possible cross-cell contamination, a minor cell-type assignment was empirically retained only when its score was within one level of the predominant cell type. For example, SCAPER was classified as rod-enriched because its rod score (“+++”) exceeded its Müller glial and endothelial scores (“+”) by two levels. Cones were not used as a primary group in this screen because cone loss in typical RP generally follows rod degeneration, although several genes are expressed in both rods and cones. Twenty-four genes were designated potentially tractable. All 24 were autosomal recessive and comprised 15 rod-specific, 3 RPE-specific, 2 Müller-glia-specific, 1 rod-RPE, and 3 RPE-Müller genes (**Figure 1C**). RPGR showed low expression in rods but higher expression in endothelial and Schwann cells and was therefore not prioritized as compatible with the present framework (**Figure S1 and Supplementary List 2**).

### Human RHO and BEST1 promoters direct cell type-enriched expression in rhesus monkey eyes

The tractable gene-selection framework depended on promoters that were active and sufficiently cell type-restricted in the primate retina. We therefore tested four human regulatory sequences previously used in rodents: RHO for rods, BEST1 for RPE, GFAP (gfaABC_1_D) for Müller glia, and RLBP1 for RPE and Müller glia (Allocca et al., 2007; Esumi et al., 2004; Lee et al., 2008; Vogel et al., 2007). Each promoter was coupled to GFP in an AAV8 vector and delivered by subretinal injection to adult male rhesus monkey eyes in ∼50uL volume. AAV8 was selected because it is publicly described and widely used and transduces ocular cell types efficiently in preclinical models (Allocca et al., 2007; Vandenberghe et al., 2011; Xiong et al., 2019; Xue et al., 2024, 2021).

In vivo fundus imaging detected GFP fluorescence at days 28 and 56 after administration of AAV8-RHO-GFP at 1 × 10^11^ or 2.5 × 10^11^ vg/eye and AAV8-RLBP1-GFP at 1 × 10^11^ vg/eye (**Figures S2 and S3**). GFP was not apparent by fundus imaging after AAV8-BEST1-GFP at 1.5 × 10^10^ or 5 × 10^10^ vg/eye or AAV8-GFAP-GFP at 1 × 10^11^ vg/eye. N was equal to one eye per condition. Both eyes of monkeys were injected, successfully creating one >5 mm diameter subretinal bleb in each treated eye. The content of vector was masked to the operator. Histological analysis at day 56 nevertheless showed GFP immunoreactivity within the photoreceptor layer after AAV8-RHO-GFP and in RPE after AAV8-BEST1-GFP (**Figure 2A**), indicating that the absence of a detectable fundus signal did not preclude RPE expression.

**Figure 2.**
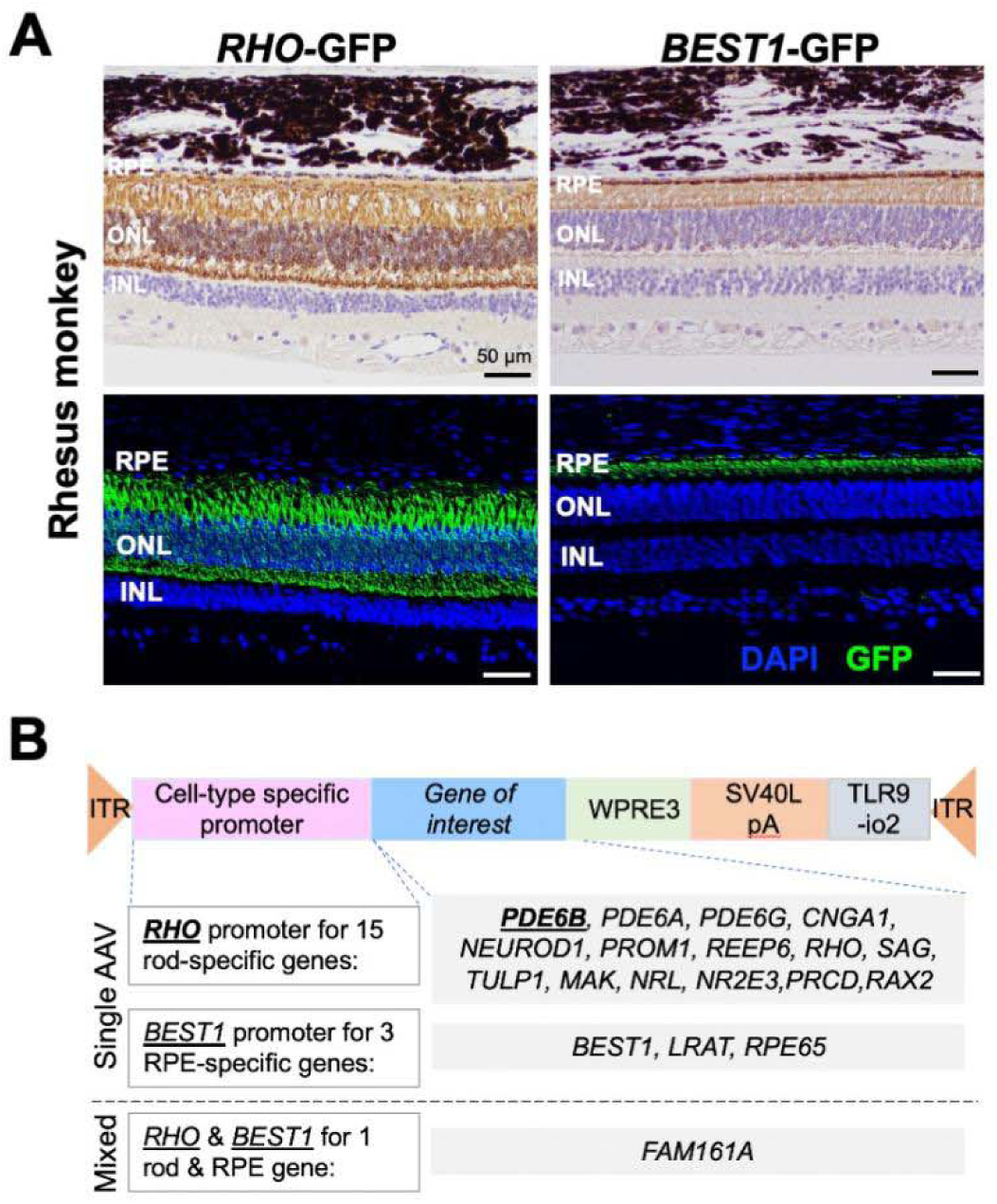
GFP expression driven by human rhodopsin (RHO) and bestrophin-1 (BEST1) promoters in rhesus monkey eyes. **(A)** Anti-GFP staining of eyes from 4-year-old male rhesus monkeys that received subretinal AAV8-RHO-GFP (2.5 × 10^11^ vg/eye) or AAV8-BEST1-GFP (1.5 × 10^10^ vg/eye). Top, H&E and DAB-HRP immunohistochemistry. Bottom, DAPI and Elab Fluor 488 immunofluorescence. **(B)** AAV vector designs for 19 autosomal recessive RP genes with predominant expression in human rods and/or RPE.

The two Müller glia-directed candidates did not provide a suitable expression profile in primate eyes. AAV8-GFAP-GFP produced no detectable GFP signal in rhesus monkey retina despite its Müller glia-specific activity in mice (**Figures S2A and S2B**). AAV8-RLBP1-GFP produced spatially heterogeneous expression together with retinal abnormalities in the monkey eye: the periphery of the subretinal bleb contained GFP-positive RPE, whereas the central region showed GFP-positive Müller glia together with RPE abnormalities (**Figure S2D**). Use of the tenfold lower mouse dose of 5 × 10^7^ vg/eye was associated with fewer RPE abnormalities but also reduced Müller glial expression (**Figure S2C**). Thus, RHO and BEST1 were retained for further platform development, whereas the GFAP and RLBP1 constructs were not.

### A shared cassette architecture supports provisional designs for 19 autosomal recessive RP genes

Combining the gene screen with the promoter results yielded 19 autosomal recessive RP genes enriched in rods and/or RPE. Fifteen genes—PDE6B, PDE6A, PDE6G, CNGA1, NEUROD1, PROM1, REEP6, RHO, SAG, TULP1, MAK, NRL, NR2E3, PRCD, and RAX2—were provisionally assigned to the human RHO promoter. BEST1, LRAT, and RPE65 were provisionally assigned to the human BEST1 promoter, and FAM161A was provisionally assigned to complementary RHO- and BEST1-driven vectors because of its expression in both rods and RPE (**Figures 1C and 2B**).

The proposed cassettes used WPRE3 together with the SV40 late polyadenylation sequence (W3SL) to conserve packaging space while supporting transgene expression (Choi et al., 2014; Loeb et al., 1999). A TLR9 inhibitory sequence, TLR9io2, was positioned adjacent to the 3′ inverted terminal repeat with the aim of reducing innate immune sensing of vector DNA (Chan et al., 2021). The resulting design therefore combined a shared AAV8 capsid and compact post-transcriptional elements with either a rod- or RPE-directed promoter.

### Dose-ranging identifies lower-dose conditions with fewer detectable retinal abnormalities in mouse and rhesus monkey eyes

We selected PDE6B as the first proof-of-concept target because it represents a relatively well-characterized patient subgroup among the 19 candidates (Kuehlewein et al., 2021), has well-defined phototransduction biology, and can be evaluated in the rapidly degenerating rd1 and rd10 mouse models (Chang et al., 2007; Deterre et al., 1988; Hartong et al., 2006; Keeler, 1924; Mclaughlin et al., 1995; Tsang et al., 2008). The therapeutic cassette consisted of AAV8-RHO-PDE6B-W3SL-TLR9io2.

Wild-type mice received subretinal injections at postnatal day 2 (P2) and were examined at P30. Eyes receiving 1.25 × 10^9^ vg/eye showed focal waviness and protrusions involving the outer nuclear layer (ONL) and inner/outer segment region. In contrast, eyes receiving 5 × 10^8^ vg/eye appeared similar to AAV-GFP-treated control eyes (**Figure 3A**). Adult rhesus monkey eyes showed a qualitatively similar dose-associated pattern. The 2.5 × 10^11^ vg/eye dose was associated with mild RPE and ONL abnormalities by optical coherence tomography (OCT) and histology, whereas fewer retinal and RPE abnormalities were detected at 1 × 10^11^ vg/eye through day 56 (**Figures 3B and S4**). These observations supported selection of 5 × 10^8^ vg/eye in neonatal mice and 1 × 10^11^ vg/eye in rhesus monkeys as the lower doses for subsequent studies under the tested conditions.

**Figure 3.**
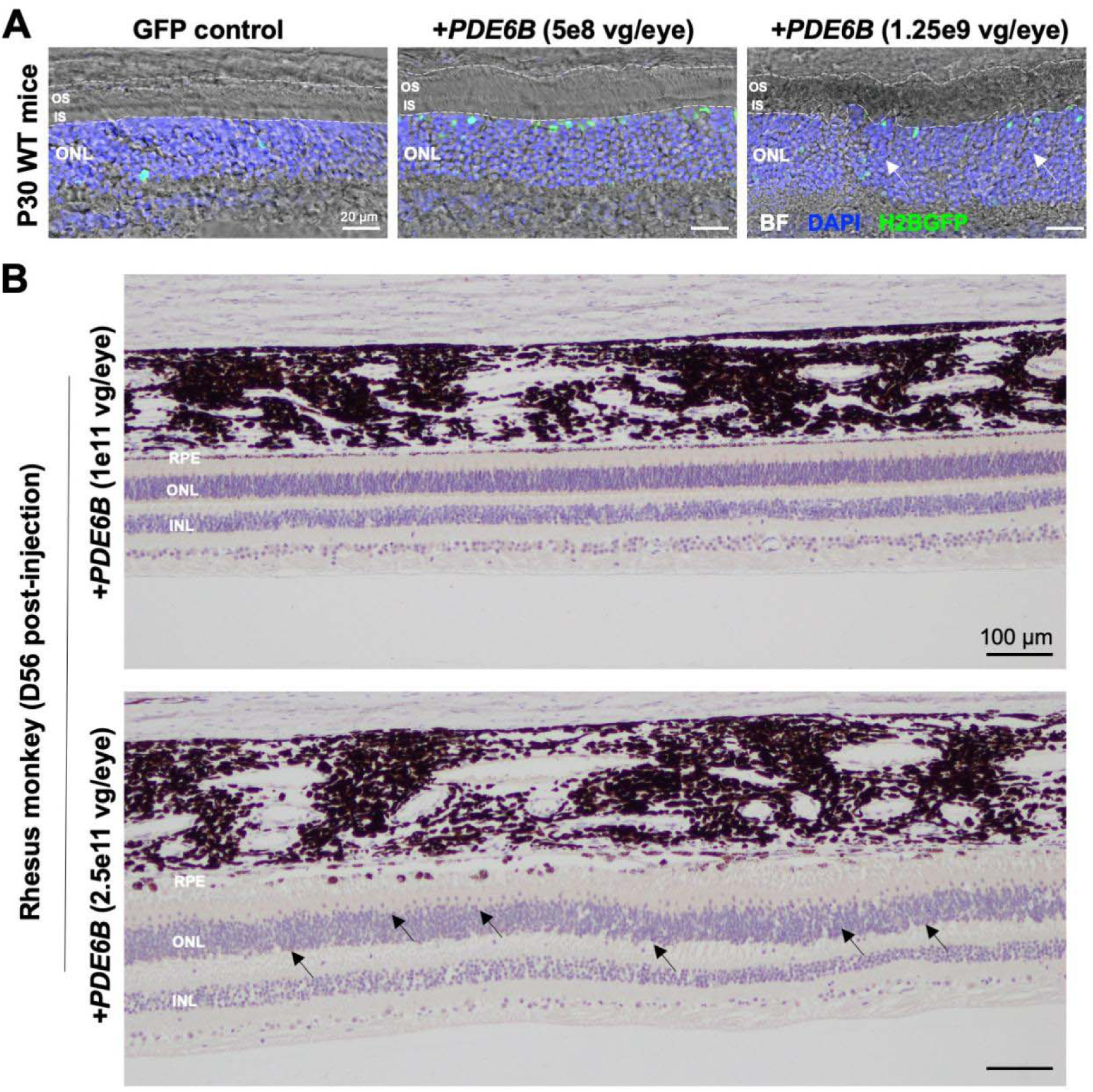
Dose-ranging and dose-associated retinal findings for AAV8-RHO-PDE6B in mouse and rhesus monkey eyes. **(A)** Representative immunofluorescence images of P30 wild-type mouse eyes after subretinal injection at P2. GFP control, 2.5 × 10^8^ vg/eye AAV8-RedO-H2BGFP. Experimental groups, 5 × 10^8^ vg/eye (low dose) or 1.25 × 10^9^ vg/eye (high dose) AAV8-RHO-PDE6B plus 2.5 × 10^8^ vg/eye AAV8-RedO-H2BGFP. Arrowheads indicate ONL and OS/IS protrusions considered potentially treatment-related. **(B)** H&E-stained sections from eyes of 4-year-old rhesus monkeys collected 56 days after subretinal injection. Top, 1 × 10^11^ vg/eye AAV8-RHO-PDE6B (low dose). Bottom, 2.5 × 10^11^ vg/eye (high dose). Arrowheads indicate disorganization of ONL nuclei.

### AAV8-RHO-PDE6B preserves retinal structure in rd1 and rd10 mice

We next tested whether the lower mouse dose delayed degeneration in two Pde6b-deficient strains with distinct disease kinetics. AAV8-RHO-PDE6B preserved ONL thickness in rd1 eyes at P20 and postnatal month 6 (PM6) and in rd10 eyes at P90 (**Figure 4A**). Longitudinal OCT imaging likewise showed greater ONL thickness in treated rd10 eyes at P30 and PM6 (**Figure 4B**). Observation of ONL preservation in both models supports that the effect was not restricted to a single Pde6b allele or rate of rod degeneration.

**Figure 4.**
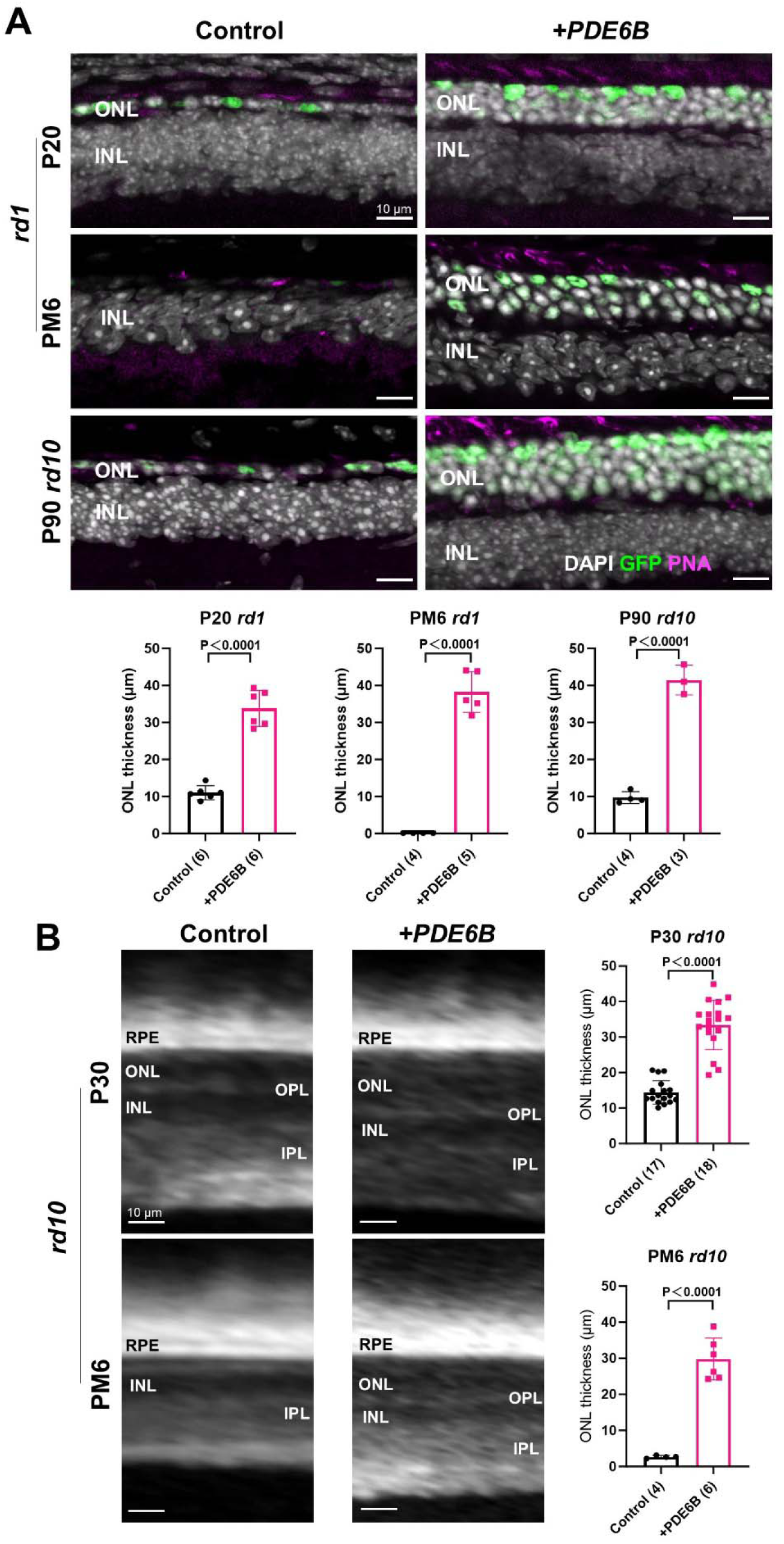
Effects of low-dose AAV8-RHO-PDE6B on photoreceptor preservation in two Pde6b-deficient mouse strains. **(A)** Representative immunofluorescence images and ONL-thickness measurements from P20 and PM6 rd1 eyes and P90 rd10 eyes after subretinal injection at P2. Control, 2.5 × 10^8^ vg/eye AAV8-RedO-H2BGFP. +PDE6B, 5 × 10^8^ vg/eye AAV8-RHO-PDE6B plus 2.5 × 10^8^ vg/eye AAV8-RedO-H2BGFP. Numbers in parentheses indicate the number of eyes analyzed. **(B)** Representative OCT images and in vivo ONL-thickness measurements from P30 and PM6 rd10 eyes after subretinal injection at P2. Treatment groups were as in (A). Data are mean ± SD; unpaired two-tailed Student’s t test.

Treatment at P0/P1 did not produce a detectable structural benefit (**Figure S5B**), which may reflect the developmental timing of rod birth and the requirement for sufficient transgene expression before degeneration (Cepko, 2014) (**Figure S5A**). As a rapid plasmid-level test of the cassette, we also electroporated pAAV-RHO-PDE6B into P1 rd1 retinas. Regions with detectable transfection showed greater ONL thickness at P20 than non-transfected regions of the same eye (**Figure S5C**), supporting expression-cassette activity independently of packaged AAV.

### AAV8-RHO-PDE6B preserves retinal responses and visually guided behavior

Electroretinography (ERG) was used to determine whether structural preservation was accompanied by retinal function. Treated rd1 eyes showed increased scotopic a- and b-wave amplitudes, consistent with improved rod and downstream rod-bipolar responses, together with increased photopic b-wave amplitudes (**Figures 5A and 5B**). Treated rd10 eyes showed a similar improvement in scotopic and photopic responses (**Figures 5C and 5D**).

**Figure 5.**
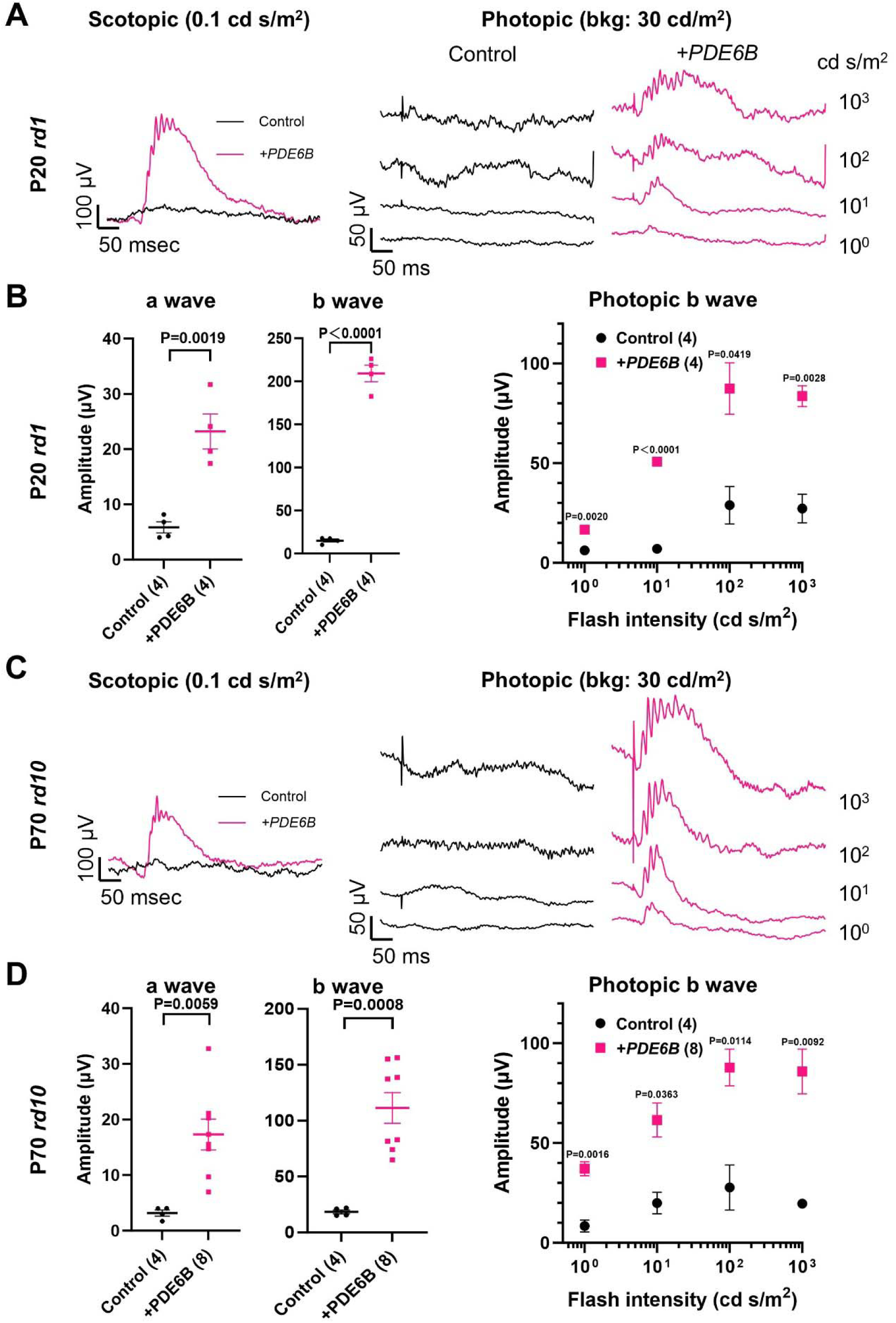
Effects of AAV8-RHO-PDE6B on retinal function in rd1 and rd10 mice. **(A)** Representative scotopic and photopic ERG responses from P20 rd1 eyes after subretinal injection at P2. Control and +PDE6B groups were as described in Figure 4. bkg, background light intensity. **(B)** Mean scotopic and photopic ERG responses from P20 rd1 eyes treated as in (A). **(C)** Representative scotopic and photopic ERG responses from P70 rd10 eyes after subretinal injection at P2. **(D)** Mean scotopic and photopic ERG responses from P70 rd10 eyes treated as in (C). Data are mean ± SEM, and analyzed using unpaired two-tailed Student’s t test with Bonferroni correction if applicable.

We then assessed cone-dependent visual behavior under photopic conditions. AAV8-RHO-PDE6B was associated with increased light avoidance in P30 rd1 mice, of which both eyes were treated with the same vector (**Figures 6A and 6B**), and preserved optomotor responses in P30 rd1 and P60 rd10 mice (**Figures 6C and 6D**). Thus, across two Pde6b-deficient models, treatment was associated with preservation of retinal structure, electrophysiological responses, and visually guided behavior.

**Figure 6.**
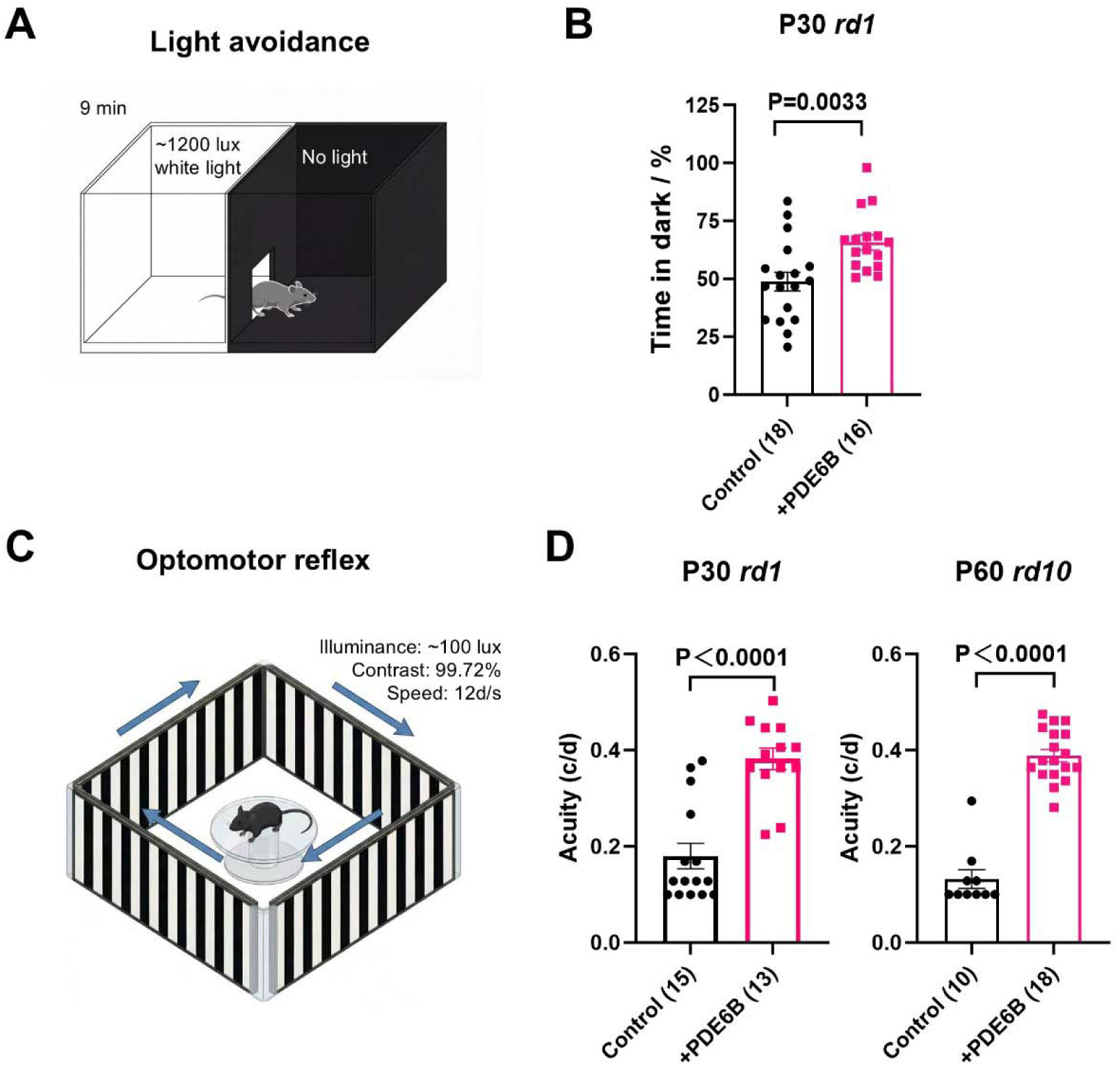
Effects of AAV8-RHO-PDE6B on visually guided behavior under photopic conditions in rd1 and rd10 mice. **(A)** Schematic of the light/dark-box assay used to measure light-dark discrimination. **(B)** Mean light-avoidance behavior of P30 rd1 mice after subretinal injection at P2. Treatment groups were as described in Figure 4. Numbers in parentheses indicate the number of analyzed animals. Data are analyzed using unpaired two-tailed Student’s t test. **(C)** Schematic of the photopic optomotor assay used to measure visual acuity. **(D)** Mean photopic optomotor responses of P30 rd1 and P60 rd10 mice after subretinal injection at P2. Eyes without a detectable response were plotted at 0.1 cycles/degree, below the 0.128 cycles/degree peak sensitivity reported for wild-type mice (Benkner et al., 2013). Treatment groups were as in (B). Numbers in parentheses indicate the number of analyzed eyes. Data are analyzed using two-tailed Mann-Whitney test. Data are mean ± SEM.

## Discussion

This study defines a selection and preclinical study framework for AAV-mediated augmentation of a genetically diverse subset of autosomal recessive RP. By integrating human retinal cell-type-specific expression, coding-sequence size, and promoter availability, we identified 19 genes compatible with a shared AAV8-based framework. Human RHO and BEST1 promoters supported rod- and RPE-enriched transgene expression, respectively, in rhesus monkey eyes. A PDE6B proof-of-concept vector was then evaluated across dose-ranging studies, two RP mouse disease models, and rhesus monkey eyes. At the lower doses tested, AAV8-RHO-PDE6B preserved retinal structure, electrophysiological responses, and visually guided behaviors in rd1 and rd10 mice, with fewer detectable abnormalities at the lower tested dose in rhesus monkey eyes through day 56. These results support further evaluation of the feasibility of reusing validated vector components across several recessive RP genotypes, but they do not establish clinical safety or efficacy for PDE6B or the other 18 genes.

The platform is intended to address a practical mismatch in IRD development. Gene augmentation is conceptually applicable to many recessive disorders, yet the patient population for each individual genotype may be too small to sustain a conventional, fully independent drug development program. A shared capsid, promoter set, cassette architecture, manufacturing process, and preclinical assay package could reduce duplicated work. The present data provide a biological basis for such reuse by linking each candidate gene to the retinal cell type in which it is predominantly expressed. However, cassette-level similarity cannot substitute for transgene-specific validation. Each product will still require confirmation of expression, potency, biodistribution, immunogenicity, and toxicology, because the encoded protein itself may alter tolerability.

### Dose, developmental timing, and promoter choice constrain translation

The dose-ranging experiments suggest a potentially narrow exposure window. Higher doses of AAV8-RHO-PDE6B were associated with retinal abnormalities in both species, whereas the lower doses were comparatively well tolerated under the conditions examined. This finding is consistent with previous evidence that AAV cis-regulatory elements and transgene expression levels contribute to ocular toxicity (Gardner et al., 2026; Xiong et al., 2019). It also argues against assuming that a cell-type-specific promoter is intrinsically safe at all vector doses or concentrations. The rhesus monkey study was limited in animal number and follow-up duration, and the absence of prominent abnormalities at 1 × 10^11^ vg/eye through day 56 should therefore be interpreted as an initial tolerated-dose observation rather than a definitive no-observed-adverse-effect level.

Treatment timing also influenced outcome. Delivery at P2 was associated with photoreceptor preservation in rd1 and rd10 mice, whereas delivery at P0/P1 did not produce a detectable benefit. This result may reflect the developmental timing of rod genesis, vector transduction, and transgene expression relative to the rapid onset of degeneration (Cepko, 2014). More broadly, it reinforces the requirement to treat while a sufficient target-cell population remains. For patients, eligibility would need to be based on genotype together with evidence of residual retinal structure and function, rather than genotype alone. Because the mouse interventions were performed neonatally, additional studies in later-stage and larger-eye models will be needed to define a clinically relevant therapeutic window.

Promoter performance did not translate uniformly from mouse to primate. The human GFAP promoter drove robust Müller glial expression in mice but did not produce detectable expression in rhesus monkey retina. The RLBP1 promoter reached Müller glia only in regions associated with RPE degeneration, and lowering the dose reduced both the retinal abnormality and Müller glial expression. These results did not support inclusion of five Müller glia-associated recessive RP genes in the current platform and emphasize the need for a primate-validated Müller glial cis-regulatory element. The observations also illustrate why promoter activity and toxicity should be tested in a relevant translational species and retinal compartment before a shared cassette is extended to additional genes (Jüttner et al., 2019).

### Relationship to previous PDE6 augmentation studies

Several groups have reported photoreceptor preservation after PDE6B augmentation in rodent and canine models (Allocca et al., 2011; Bennett et al., 1996; Jomary et al., 1997; Pang et al., 2008; Petit et al., 2012; Pichard et al., 2016). The HORA-PDE6B-001 program used an AAV5-GRK1-PDE6B vector and was recorded as terminated on December 18, 2025, because of lack of financing rather than a reported biological failure (ClinicalTrials.gov #NCT03328130). The present vector differs in capsid, promoter, compact post-transcriptional elements, and inclusion of TLR9io2. The human RHO promoter is more restricted to rods than GRK1 in primate retina, where GRK1 can also drive cone expression (Boye et al., 2012). Nevertheless, differences among studies in capsid, promoter, titer, delivered volume, animal model, and manufacturing prevent attribution of performance to any single element.

The recent PDE6A phase I/IIa study is an important caution for platform development. Despite supportive preclinical studies, a nine-participant trial of AAV8-RHO-PDE6A did not improve visual function over 1 year and reported persistent ocular findings, including retinal thinning and visual-acuity loss in some participants (Reichel et al., 2026; Schön et al., 2017; Seitz et al., 2025; Wert et al., 2013). Although the PDE6A cassette differed from the present design in its post-transcriptional and innate immune-modulatory elements, these differences do not establish that W3SL or TLR9io2 will improve clinical outcomes. Larger preclinical cohorts, longer follow-up, clinically matched delivery parameters, and transgene-specific toxicology will be necessary before advancing PDE6B or another platform gene.

### A staged access model for rare, early-stage genotypes

We outline a not-for-profit development model in which genetically confirmed patients with one of the candidate genes would undergo structural assessment of the peripheral ONL and assessment of pre-existing AAV8 immunity using a prospectively validated assay and cutoff, if clinically justified (**Figure 7**). Of note, these three criteria are not the only ones to screen eligible patients for enrollment. Initial administration would occur only within an appropriately authorized clinical study (e.g. under the regulation of National Health Commission, NHC, in China taken effect on May 1^st^, 2026), with safety and preservation of remaining retinal structure and function evaluated over time. For early-stage RP, outcomes based solely on best-corrected visual acuity may be insensitive because central cone-mediated acuity can remain preserved while peripheral rods are lost. Structural measures, visual fields, and rod- and cone-pathway functional tests may therefore be needed, with the final endpoint set prospectively defined for each genotype and specific disease stage.

**Figure 7.**
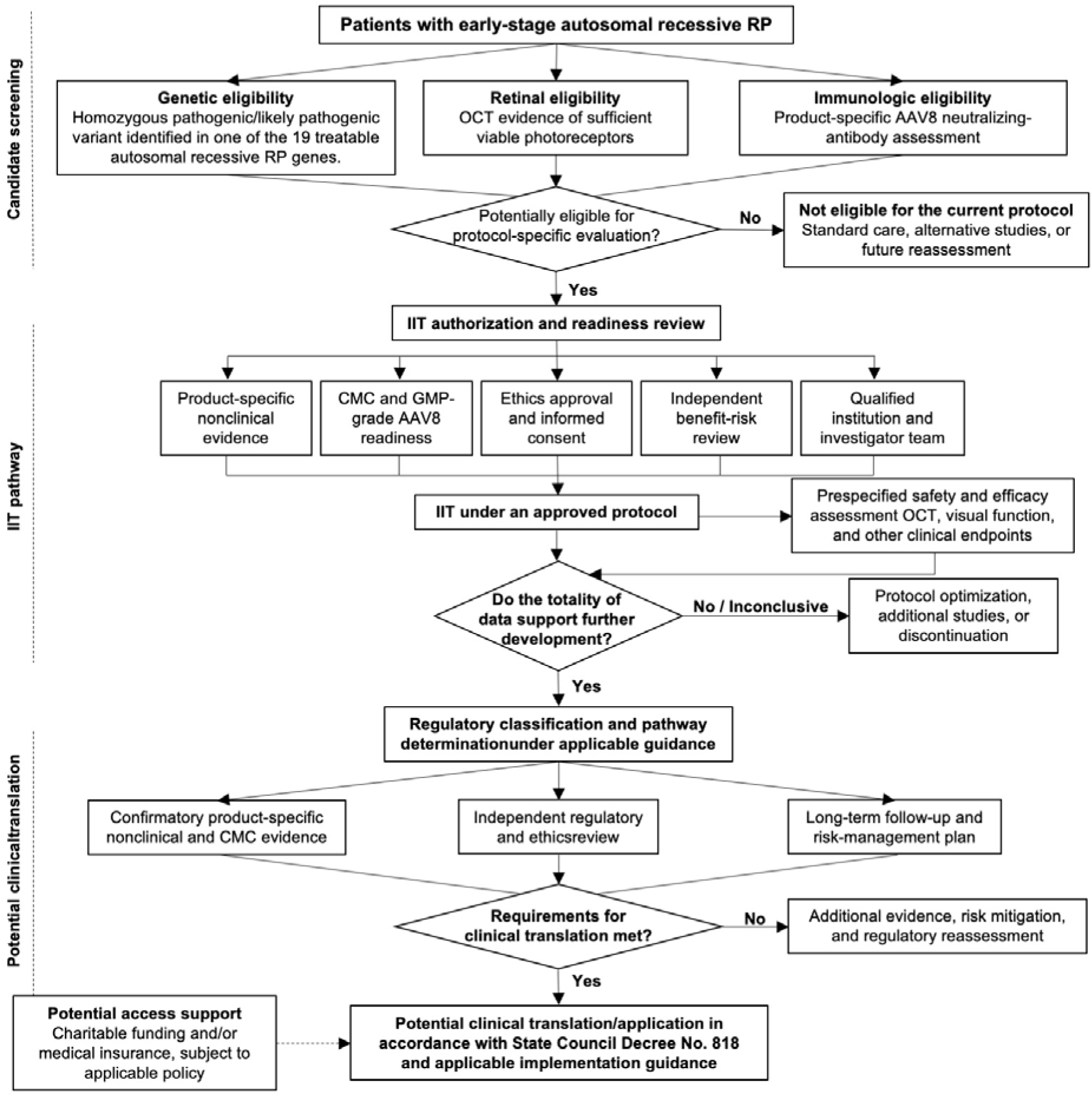
Conceptual patient-selection, IIT, and Clinical-Translation Pathway for AAV8 gene therapy in RP. RP, retinitis pigmentosa; OCT, optical coherence tomography; ONL, outer nuclear layer; IIT, investigator-initiated trial; GMP, good manufacturing practice; CDMO, contract development and manufacturing organization; NHC, National Health Commission; State Council Decree No. 818, Regulations on the Administration of Clinical Research and Clinical Translational Application of New Biomedical Technologies.

This proposed access pathway is a development concept rather than a clinical protocol. Regulatory classification, manufacturing standards, trial authorization, long-term follow-up, pharmacovigilance, liability and insurance, and sustainable funding would need to be established with hospitals, regulators, manufacturers, patients, and independent ethics committees. The framework may lower repeated preclinical and operational costs, but it cannot remove the requirement for product-specific evidence. In China, the final gene therapy product either falls under the drug regulation of National Medical Products Administration, or NHC pathway that could be provided at a few designated hospitals.

### Limitations and future directions

Several limitations define the next experiments. First, the gene-selection screen relies on existing single-cell datasets and semiquantitative expression thresholds; low-abundance transcripts, regional expression, developmental changes, and dataset-specific dropout may affect classification. Second, promoter evaluation and PDE6B ocular tolerability testing in rhesus monkeys used small cohorts and a 56-day endpoint. Third, efficacy was established in neonatal mouse models with rapid degeneration, not in adult large-animal disease models. Fourth, the work does not yet report validated vector-genome titration, full/empty capsid characterization, formal biodistribution, immunogenicity, good laboratory practice toxicology, or a manufacturing comparability program. Finally, the remaining 18 candidate transgenes were assigned by biological compatibility but were not individually tested.

Future work should therefore validate each cassette experimentally, expand non-human primate dose and duration studies, develop a primate-active Müller glial promoter, and define genotype-appropriate clinical endpoints. Parallel approaches will still be needed for dominant disease (Pulman et al., 2022) (**Figure S1**), genes exceeding AAV packaging capacity (Ghosh et al., 2011), late-stage degeneration (Lüscher et al., 2025; Xue and Cepko, 2023), disorders requiring broad retinal delivery (Agrawal et al., 2025), and non-AAV-based approaches (Yao et al., 2025; Zeng et al., 2025). Within these boundaries, the present study provides a proposed starting point for prioritizing recessive RP genes and testing shared components without conflating platform efficiency with product-level validation.

## Conclusion

By integrating human retinal cell-type expression, AAV packaging constraints, and promoter performance in rhesus monkeys, this study provisionally identifies 19 autosomal recessive RP genes that are candidates for a shared gene-augmentation framework. The PDE6B proof-of-concept data support structural and functional rescue in two mouse models, but the limited primate cohort, short follow-up, neonatal treatment paradigm, and absence of product-specific testing for the other 18 genes preclude clinical conclusions. The framework therefore provides a prioritized starting point for transgene-specific validation rather than a substitute for vector-level safety, efficacy, manufacturing, and regulatory studies.

## Materials and methods

### Animals

We used two wild-type mouse strains verified to lack the rd8 mutation (Mattapallil et al., 2012): C57BL/6JNifdc (Vital River Beijing, stock 219) and Crl:CD1(ICR) (Vital River Beijing, stock 201). Pde6b^rd1^ mice (MGI:1856373) were maintained on FVB/NCrl (Charles River, stock 207), C3H/HeJ (Jackson Laboratory, stock 000659), or C57BL/6J backgrounds (gift from Connie Cepko, Harvard University) (Chandler et al., 2026). Pde6b^rd10^ mice (MGI:2388259) were maintained on a C57BL/6J background (gift from Ying Xu, Jinan University; originally from Jackson Laboratory, stock 004297). Mice were housed under specific-pathogen-free conditions on a 12-h light/12-h dark cycle at 22 ± 2°C and 50% ± 10% humidity, with food and water available ad libitum. Both male and female mice were randomized and used without distinguish in all experiments. After weaning, male and female mice were housed separately till the experimental end points. Both eyes were injected and used in all experiments. All mouse procedures were performed at Lingang Laboratory and approved by its Institutional Animal Care and Use Committee (IACUC; protocol NZXSP-2022-4). All monkey procedures were performed at PriMed facilities accredited by AAALAC International and were approved by the PriMed IACUC (protocol AW2520). The monkeys were quarantined and acclimated for 4 weeks before experimentation and were screened for serum AAV8-neutralizing antibodies (Day et al., 2018). Only antibody-negative animals were included. Eight eyes from five healthy male rhesus monkeys 4–6 years of age were used for this study. Both eyes were treated with different vectors or dosages. Two eyes were unreported due to irrelevant vector content to this study. Monkeys were housed at 18°C–26°C and 40%–70% humidity on a 12-h light/12-h dark cycle, with at least eight air exchanges per hour using 100% fresh air.

### Plasmid construction

Plasmids were constructed by Gibson assembly as described previously (Gibson et al., 2009; Rabe and Cepko, 2020; Xue et al., 2024, 2021). DNA fragments were amplified from existing cDNA templates or synthesized commercially (Genewiz, Suzhou, China). Components included AAV2 ITRs, the human RHO promoter (−800 to +6 bp fragment) (Allocca et al., 2007), the human BEST1 promoter (−540 to -2 bp fragment) (Esumi et al., 2004; Wu et al., 2021), the human GFAP promoter (the 681 bp gfaABC_1_D; gift from Wenjun Xiong, City University of Hong Kong) (Lee et al., 2008; Wu et al., 2025), WPRE3 and SV40 late polyadenylation sequences (Choi et al., 2014), and TLR9io2 (Chan et al., 2021). A human RLBP1 promoter (−3376 to −2283 bp fragment, National Center for Biotechnology Information locus ID 6017) was synthesized and cloned based on the reported locus (Vogel et al., 2007) and compared with a mouse Rlbp1 promoter construct (Matsuda and Cepko, 2004). Human cDNAs for the 19 autosomal recessive RP genes were obtained from NCBI records (for example, PDE6B, NM_000283.4) and synthesized commercially. CAG-GFP was provided by the Cepko laboratory (Matsuda and Cepko, 2004). Plasmids were amplified in Stbl3-competent Escherichia coli and validated by XmaI digestion to assess ITR integrity, Sanger sequencing across assembly junctions, and complete plasmid sequencing (Genewiz, Suzhou, China). All AAV genome sizes (ITR-to-ITR) were smaller than 4.8 kb. Plasmid sequence maps are available upon request.

### AAV preparation

Vectors were packaged in AAV-293 cells with AAV8 capsids and purified by iodixanol-gradient ultracentrifugation in a biosafety level 2 facility at Lingang Laboratory, as described previously (Grieger et al., 2006; Xiong et al., 2015; Xue et al., 2021). Titers were estimated by protein gel electrophoresis against serially diluted AAV standards, which were prepared in the same way with previously established qPCR titration and in vivo infectious efficiency. The full/empty ratio of capsid was not examined in this pre-clinical exploratory study. Vector concentrations ranged from 2 × 10^12^ to 3 × 10^13^ vg/mL. Empirically, this titration method sufficiently provided consistent and reliable in vivo preclinical outcomes.

### Histology

Mice were euthanized by cervical dislocation, and eyes were immediately enucleated and fixed in 4% paraformaldehyde for 2 h at room temperature with gentle rocking (Xue et al., 2024, 2021). After removal of the cornea, lens, and iris, eyecups were dehydrated, embedded, and cryosectioned at 20 μm. Sections were stained with DAPI and peanut agglutinin conjugated to Cy5 or rhodamine (1:1,000) to visualize nuclei and the cone extracellular matrix, respectively. Confocal images were acquired on Olympus FLUOVIEW FV3000 or Leica STELLARIS 5 systems using 40×/0.95 NA air objectives. All sections were examined by the researcher, and one representative field per mouse eye was presented in this study. The mouse pathologist was not masked to the experimental conditions. Monkey eyes were collected on day 56 after completion of ophthalmic examinations. Animals were deeply anesthetized by intravenous ketamine/xylazine (30/20 mg/kg), euthanized by femoral-artery exsanguination, and immediately enucleated. Eyes were fixed in formalin-acetic acid-alcohol for at least 24 h, opened at the limbus, and dissected to remove the anterior segment and vitreous. Tissue from the injection site was dehydrated, paraffin-embedded, and sectioned. For immunohistochemistry, sections were deparaffinized, rehydrated, subjected to EDTA antigen retrieval, and incubated with primary anti-GFP antibody (ab290) overnight at 4°C. After incubation with an HRP-conjugated secondary antibody polymer, signal was developed with DAB and sections were counterstained with hematoxylin. For immunofluorescence, rehydrated sections were blocked with goat serum (E-IR-R110), incubated with anti-GFP antibody overnight at 4°C, and labeled with Elab Fluor 488-conjugated goat anti-rabbit IgG (E-AB-1055) for 60 min at 37°C before DAPI staining. Immunohistochemical images were acquired using an Olympus BX43F microscope, and immunofluorescence images were acquired using a 3DHISTECH Pannoramic Scan digital pathology scanner. All sections were examined by the researcher, and one representative field per monkey eye was presented in this study. The monkey pathologist was masked to the experimental conditions.

### Subretinal injection

For mice, AAV vectors or plasmids were randomized and injected subretinally into neonatal eyes as described previously (Matsuda and Cepko, 2004; Xiong et al., 2015). The mouse researcher was not masked to the vector identity. For monkeys, operators were blinded to vector identity until the manuscript preparation phase. Monkeys were anesthetized by intramuscular ketamine/dexmedetomidine (20/0.015 mg/kg), pupils were dilated with tropicamide, and the periocular skin and ocular surface were disinfected with povidone-iodine. A sclerotomy was made 2–4 mm posterior to the limbus, and a 40-gauge beveled cannula was advanced through the scleral tunnel into the vitreous cavity. Under fundus visualization, vector suspended in 50 μL PBS was delivered subretinally within the superior vascular arcade while avoiding retinal vessels, creating a single > 5 mm diameter bleb which was confirmed by post-operational OCT and fundus imaging. No steroids were co-injected with the AAVs. Fundus photography and OCT were performed immediately after injection, usually showing a >5 mm diameter subretinal bleb. Animals were monitored during recovery and received tobramycin-dexamethasone eye drops in both eyes three times daily for 3 days.

### Fundus imaging and optical coherence tomography (OCT)

For mice, scanning laser ophthalmoscopy (SLO) and OCT were performed using the Saris non-invasive in vivo imaging platform (Robotrak, Nanjing, China), similar to previous studies (Xiong et al., 2019). The sampling location of mouse OCT was collected at representative GFP positive region, which indicated well infected area that could be visualized via the GFP channel of SLO images, in this study. In monkeys, fundus photography was performed using a Topcon TRC-50DX camera before injection, immediately after injection, and on days 28 and 56. Animals were anesthetized, and pupils were dilated as described above. At least seven white-light images were acquired per eye, centered on the macula, optic disc, superior and inferior macular branch vessels, superonasal and inferonasal optic-disc regions, and temporal macula. Green-fluorescence images were acquired using the fluorescein angiography mode and centered on the injection site to detect GFP. Imaging angle and exposure were attempted to hold consistent for each region across sessions. OCT was performed during the same sessions using a Heidelberg SPECTRALIS system. The sampling location of monkey OCT was marked on the fundus image presented in this study.

### Electroretinography (ERG) in mice

ERGs were recorded from C57BL/6J.Pde6brd1 and rd10 mice using an Espion E3 system (Diagnosys) as described previously (Avilés et al., 2025; Xue et al., 2023). Photopic responses were recorded on a 30 cd/m² background. Scotopic and photopic a- and b-wave amplitudes were quantified to assess retinal function.

### Visually guided behavioral assays in mice

Photopic optomotor acuity was measured in C57BL/6J.Pde6b^rd1^ and rd10 mice, of which both eyes received identical content of injection, using an OptoDrum system (Striatech, Germany) under approximately 100 cd/m² background illumination. Stripe contrast was 99.72%, and rotation speed was 12°/s. Visual acuity, defined as the highest spatial frequency eliciting a response, was determined automatically using a published algorithm (Benkner et al., 2013). Eyes that did not respond at 0.128 cycles/degree were assigned a value of 0.1 cycles/degree for plotting and analysis. Responses from two eyes were distinguished and recorded independently based on different preference of grating rotation direction (Douglas et al., 2005).

### Light-dark discrimination assay in mice

Light-avoidance behavior was measured in P30 C3H/HeJ and C57BL/6J.Pde6brd1 mice,, of which both eyes received the same content of injection, using an LE816 automated light/dark box with PPCWIN software (Panlab, Barcelona, Spain), as described previously (Wang et al., 2020). The chamber (51.5 × 26.5 × 27 cm) comprised equally sized dark and illuminated compartments; the illuminated compartment was maintained at approximately 1,200 lux with white LED light. The temperature difference between compartments was <1°C. Each mouse was placed in the illuminated compartment and tracked for 9 min. Time in each compartment was measured using floor-mounted piezoelectric sensors, and the percentage of total time spent in the dark compartment was calculated.

### Statistics

For ONL histology, OCT, ERG and optomotor assays, two eyes from each animal were analyzed as independent samples. For OCT and ONL histology, only one representative image from infected area per eye was used for statistical analysis. Two-tailed unpaired student’s t test or Mann-Whitney test was used for the statistical analysis via Microsoft Excel or Prism softwares. For ERG analyses, P values were adjusted for multiple comparisons using the Bonferroni correction. P or adjusted P value of < 0.05 was considered statistically significant.

## Acknowledgements

We thank members and organizers of the RP Light Love Alliance, a patient-led non-profit organization in China, for discussions; Connie Cepko (Harvard Medical School), Wenjun Xiong (City University of Hong Kong), and Ying Xu (Jinan University) for materials and animals; and Meng Zhu (University of Cambridge) for comments on the manuscript. We also thank Yizhou Jiang of Xue Lab, the equipment platforms at Lingang Laboratory and the Optical Imaging Core Facility of the Shanghai Research Center for Brain Science and Brain-Inspired Intelligence for technical support. This work was supported by Lingang Laboratory (Grant No. LGL-6666).

## Declaration of generative AI and AI-assisted technologies in the manuscript preparation process

During preparation of this work, the authors used the Xue Lab Agent built on OpenClaw (powered by ChatGPT) to polish and restructure the final version of the manuscript, including Figure 7, and used Nano Banana Pro (powered by Gemini) to create the schematic illustrations in Figures 6A and 6C. The authors reviewed and edited all outputs as needed and take full responsibility for the content of the manuscript.

## Supplementary Material

**Figure S1.**
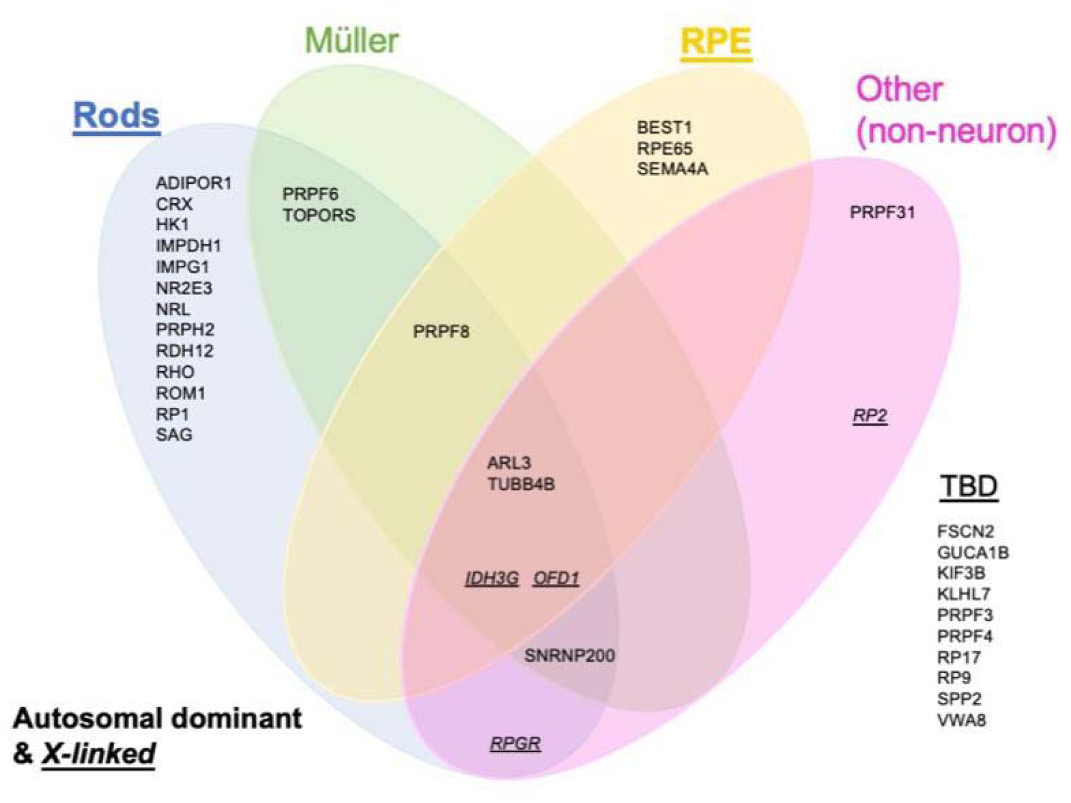
Classification of autosomal dominant and X-linked RP genes. X- linked genes are italicized and underlined.

**Figure S2.**
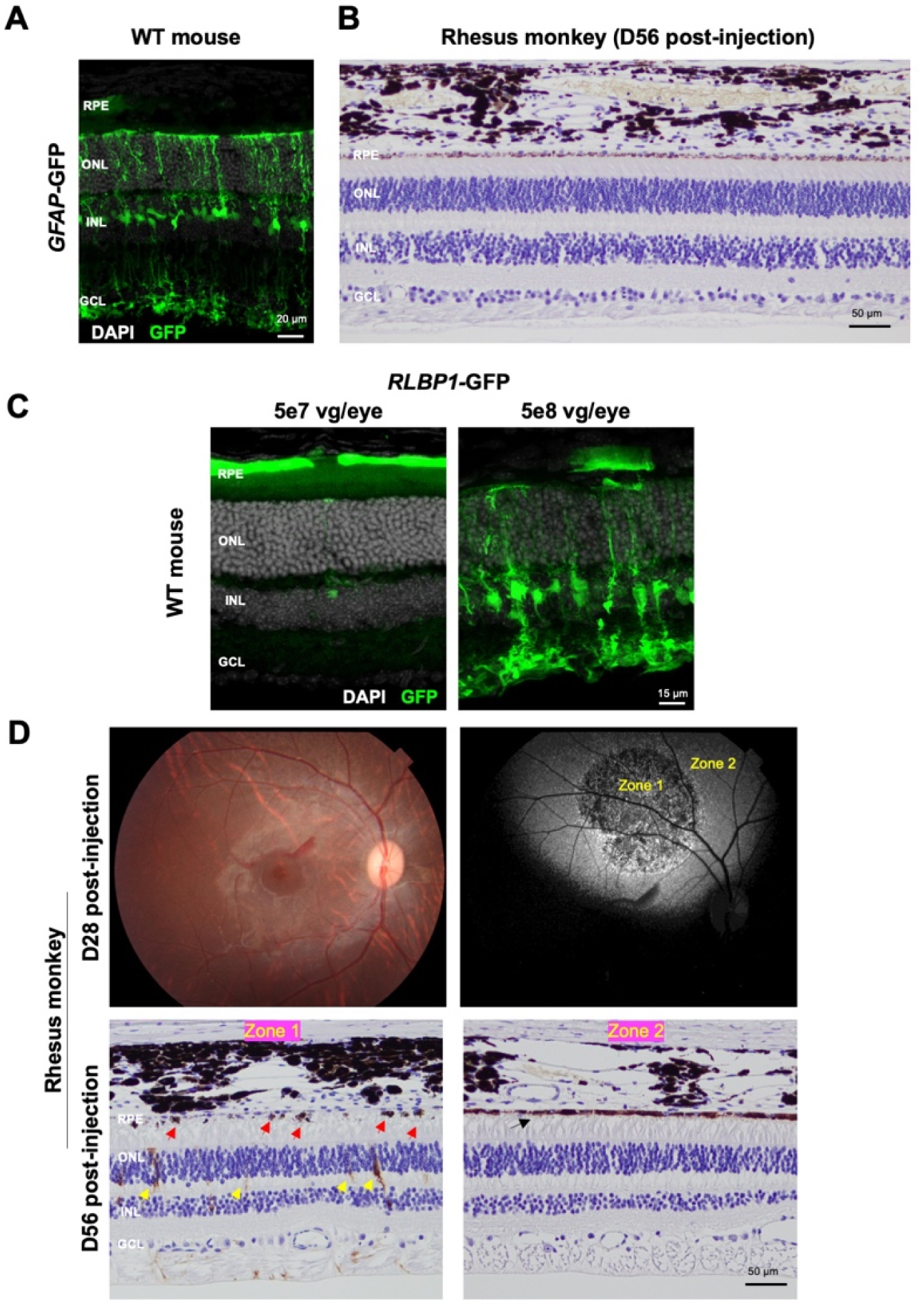
GFP expression driven by human GFAP and RLBP1 promoters in mouse and rhesus monkey eyes. **(A)** Representative cross-sectional fluorescence image of a P30 wild-type mouse eye after subretinal injection of 5 × 10^8^ vg/eye AAV8-GFAP-GFP at P4. **(B)** Representative immunohistochemical section from an eye of a 5- to 6-year-old male rhesus monkey after subretinal injection of 1 × 10^11^ vg/eye AAV8-GFAP-GFP. **(C)** Representative cross-sectional fluorescence images of wild-type mouse eyes after subretinal injection of 5 × 10^7^ vg/eye (P30) or 5 × 10^8^ vg/eye (P22) AAV8-RLBP1-GFP at P4. **(D)** White-light and fluorescence fundus images from an eye of a 5- to 6-year-old male rhesus monkey after subretinal injection of 1 × 10^11^ vg/eye AAV8-RLBP1-GFP. Zone 1, region with greater retinal abnormalities; zone 2, region with fewer retinal abnormalities. Red arrowhead, RPE abnormality; yellow arrowhead, GFP-positive Müller glia; black arrowhead, GFP-positive RPE.

**Figure S3.**
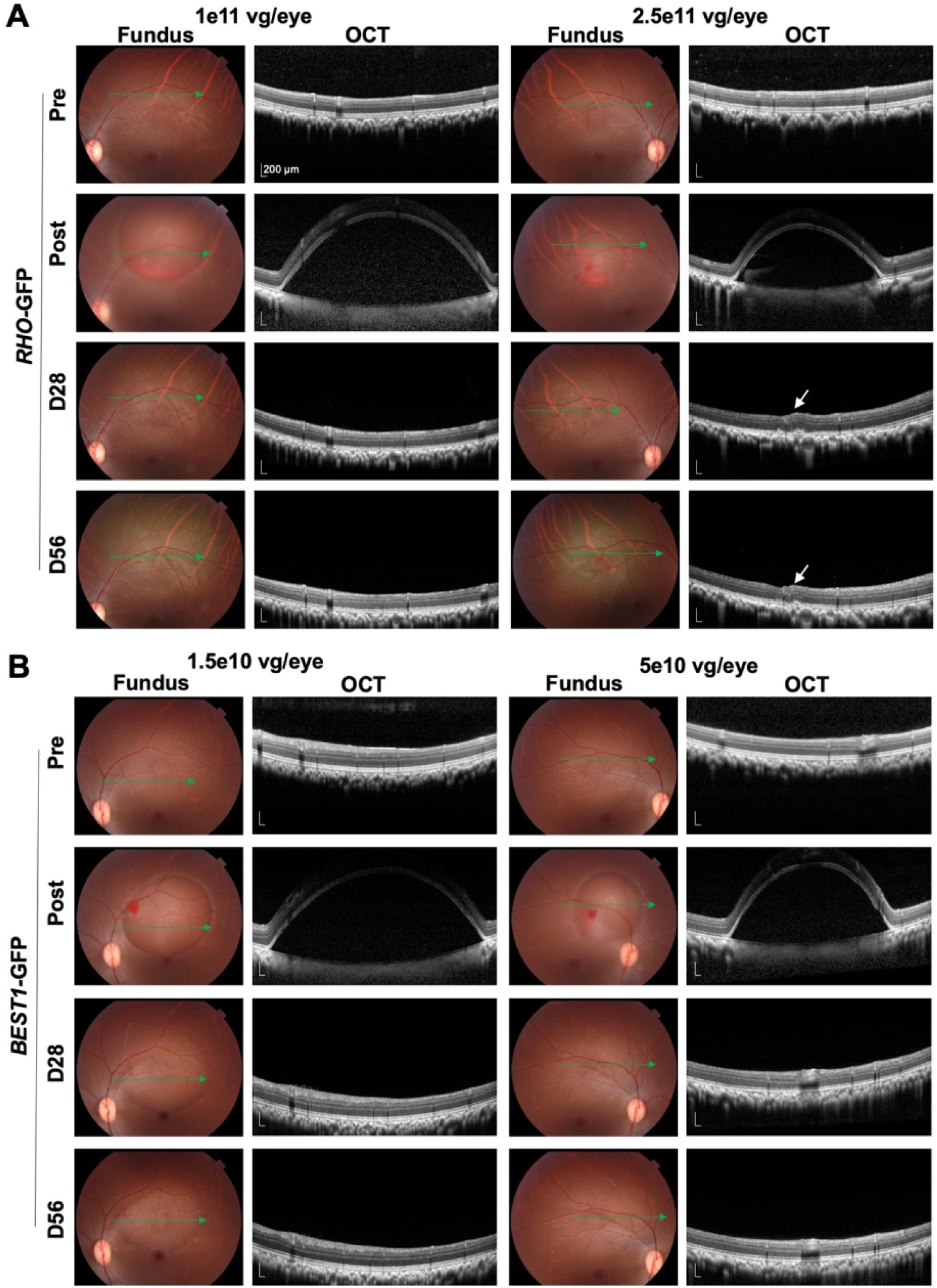
Dose-associated ocular findings after AAV8-RHO-GFP and AAV8-BEST1-GFP in rhesus monkey eyes. **(A)** White-light fundus and OCT images from eyes of 4-year-old male rhesus monkeys after subretinal injection of 1 × 10^11^ vg/eye (low dose) or 2.5 × 10^11^ vg/eye (high dose) AAV8-RHO-GFP. Green arrows indicate the OCT scan location; white arrowheads indicate retinal abnormalities. **(B)** White-light fundus and OCT images after subretinal injection of 1.5 × 10^10^ or 5 × 10^10^ vg/eye AAV8-BEST1-GFP. Green arrows indicate the OCT scan location; white arrowheads indicate regional retinal abnormalities. Pre, before injection; Post, immediately after injection; D28 and D56, 28 and 56 days after injection, respectively.

**Figure S4.**
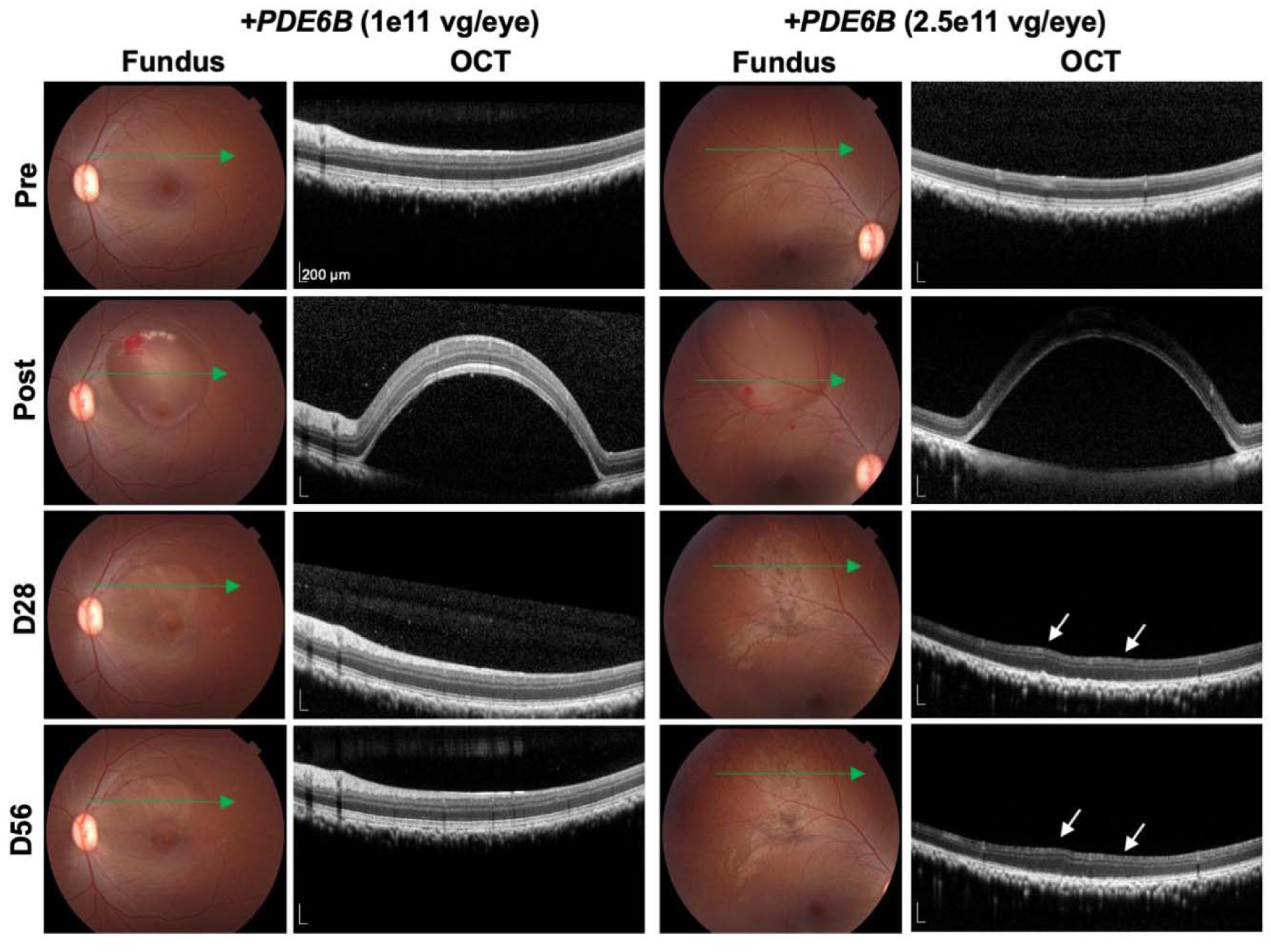
Dose-associated ocular findings after AAV8-RHO-PDE6B in rhesus monkey eyes. White-light fundus and OCT images from eyes of 4-year-old male rhesus monkeys after subretinal injection of 1 × 10^11^ vg/eye (low dose) or 2.5 × 10^11^ vg/eye (high dose) AAV8-RHO-PDE6B. Green arrows indicate the OCT scan location; white arrowheads indicate retinal abnormalities.

**Figure S5.**
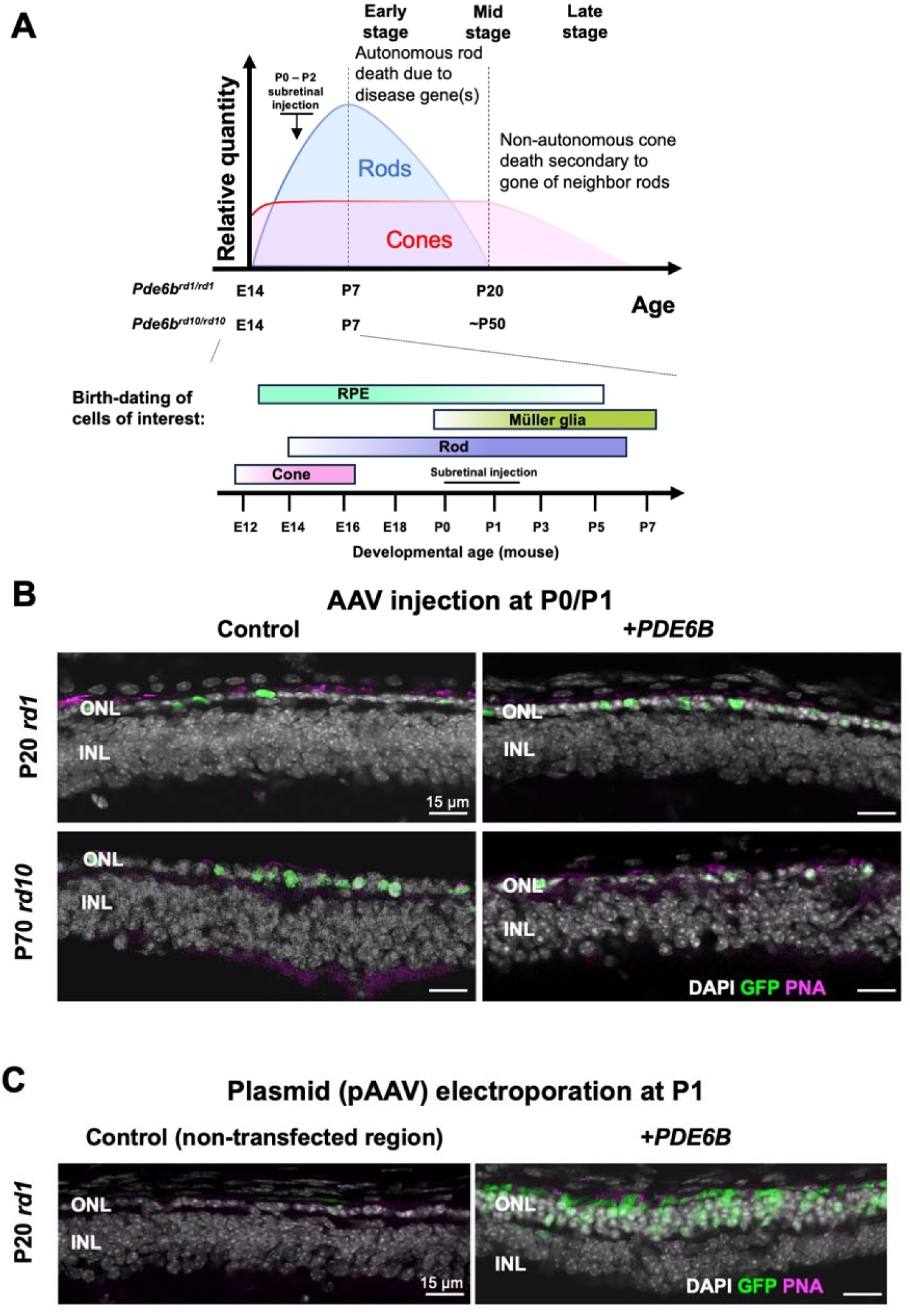
Effects of intervention timing relative to rod birth. **(A)** Schematic of retinal cell birth and photoreceptor degeneration in RP mice. (Bodenstein and Sidman, 1987; Cepko, 2014) **(B)** Representative immunofluorescence images showing ONL thickness in P20 rd1 (top) and P70 rd10 (bottom) eyes after subretinal injection at P0/P1. Control and +PDE6B groups were as described in Figure 4. **(C)** Representative immunofluorescence images showing ONL thickness in P20 rd1 eyes after subretinal injection and electroporation of AAV plasmids at P1. Control, non-transfected region of the same eye; +PDE6B, 2,000 ng/μL pAAV-RHO-PDE6B plus 300 ng/μL CAG-GFP.

